# MTL functional connectivity predicts stimulation-induced theta power

**DOI:** 10.1101/320663

**Authors:** E. A. Solomon, R. Gross, B. Lega, M. R. Sperling, G. Worrell, S. A. Sheth, K. A. Zaghloul, B. C. Jobst, J. M. Stein, S. Das, R. Gorniak, C. Inman, S. Seger, J. E. Kragel, D. S. Rizzuto, M. J. Kahana

**Affiliations:** University of Pennsylvania, Department of Bioengineering, Philadelphia, PA 19146; Department of Neurosurgery, Emory School of Medicine, Atlanta, GA 30322; Department of Neurosurgery, University of Texas Southwestern, Dallas, TX 75390; Department of Neurology, Thomas Jefferson University Hospital, Philadelphia, PA 19107; Department of Neurology, Department of Physiology and Bioengineering, Mayo Clinic, Rochester, MN 55905; Department of Neurosurgery, Baylor College of Medicine, Houston, TX 77030; Surgical Neurology Branch, National Institutes of Health, Bethesda, MD 20814; Department of Neurology, Dartmouth Medical Center, Lebanon, NH 03756; Department of Radiology, Hospital of the University of Pennsylvania, Philadelphia, PA 19104; Department of Radiology, Thomas Jefferson University Hospital, Philadelphia, PA 19107; University of Pennsylvania, Department of Psychology, Philadelphia, PA 19146

## Abstract

Focal electrical stimulation of the brain incites a cascade of neural activity that propagates from the stimulated region to both nearby and remote areas, offering the potential to control the activity of brain networks. Understanding how exogenous electrical signals perturb such networks in humans is key to its clinical translation. To investigate this, we applied electrical stimulation to subregions of the medial temporal lobe in 26 neurosurgical patients fitted with indwelling electrodes. Networks of low-frequency (5-13 Hz) spectral coherence predicted stimulation-evoked changes in theta (5-8 Hz) power, but only when stimulation was applied in or adjacent to white matter. Furthermore, these power changes aligned with control-theoretic predictions of how exogenous stimulation flows through complex networks, such as a dispersal of induced activity when functional hubs are targeted. Our results demonstrate that functional connectivity is predictive of causal changes in the brain, but that access to structural connections is necessary to observe such effects.

## Introduction

Intracranial brain stimulation is increasingly used to study disorders of human behavior and cognition, but very little is known about how these stimulation events affect neural activity. Though several recent studies have demonstrated the ability to modulate human memory with direct electrical stimulation (DES) of the cortex ^1–7^, none have described the mechanism by which stimulation yields altered cognitive states. However, understanding how the brain responds to these exogenous currents is necessary to ultimately develop therapeutic interventions that rely on DES ^8,9^.

Because it is often not possible to directly stimulate a given brain region of interest in clinical populations of neurosurgical volunteers, recent investigations have asked whether the brain’s intrinsic functional or anatomical architecture can predict how mesoscale stimulation events propagate through the brain. In monkeys, Logothetis, et al. (2010) demonstrated that the effects of electrical stimulation propagated through known anatomical connections in the visual system. In humans, corticocortical evoked potentials (CCEPs), measured with intracranial EEG, have also been shown to propagate through anatomical and functional connections ^10,11^, as has the fMRI BOLD response to stimulation ^12^. These studies provide powerful evidence that the effects of stimulation are determined by the connectivity profile of a targeted region. More broadly, renewed interest in the idea of the brain as a controllable network ^13–15^ raises a testable hypothesis in need of empirical validation: to what extent does a brain’s network architecture predict the cascade of physiologic change that accompanies a stimulation event?

In this study, we asked whether the functional connectivity of a stimulated region predicts where we observe changes in neural activity. To expand on prior work that has examined network architecture and stimulation, we adopted a paradigm that (1) measures stimulation’s effect on low-frequency (theta) power, a cognitively-relevant electrophysiological biomarker, and (2) simultaneously considers the structural and functional connectivity of a targeted region.

In 26 neurosurgical patients with indwelling electrodes, we stimulated different regions of the medial temporal lobe (MTL) and asked whether functional connectivity predicted modulations of theta power in distributed cortical regions. We showed that functional connectivity was only predictive of theta modulation when stimulation occurred in or near a white matter tract, but in those cases, stimulation could evoke sustained increases in theta power even in distant regions. Furthermore, functional networks only had such predictive power at low frequencies, in the theta and alpha bands (5-13 Hz).

## Results

### Calculating a theta modulation index

To determine how direct cortical stimulation propagates through brain networks, we collected intracranial EEG (iEEG) data from 26 patients undergoing clinical monitoring for seizures. Subjects rested passively in their hospital bed while we applied bipolar macroelectrode stimulation at varying frequencies (10-200 Hz) and amplitudes (0.25 to 1.5 mA) to MTL depth electrodes (see online Methods for details). Rectangular stimulation pulses were delivered for 500 ms, followed by a 3-second inter-stimulation interval (Figure 1A-C). Each subject received at least 240 stimulation events (“trials”) at 1-8 distinct sites in MTL gray or white matter (mean 2.7 sites; see Supplementary Table 2 for stimulation locations). During a separate recording session in which no stimulation occurred, for each subject we computed resting networks of low-frequency (5-13 Hz) coherence, motivated by prior literature that shows robust iEEG functional connectivity at low frequencies ^16–19^. These networks reflect correlated low-frequency activity between all possible pairs of electrodes in a subject, during a period when subjects are passively waiting for a task to begin (Figure 2A).

**Figure 1.**
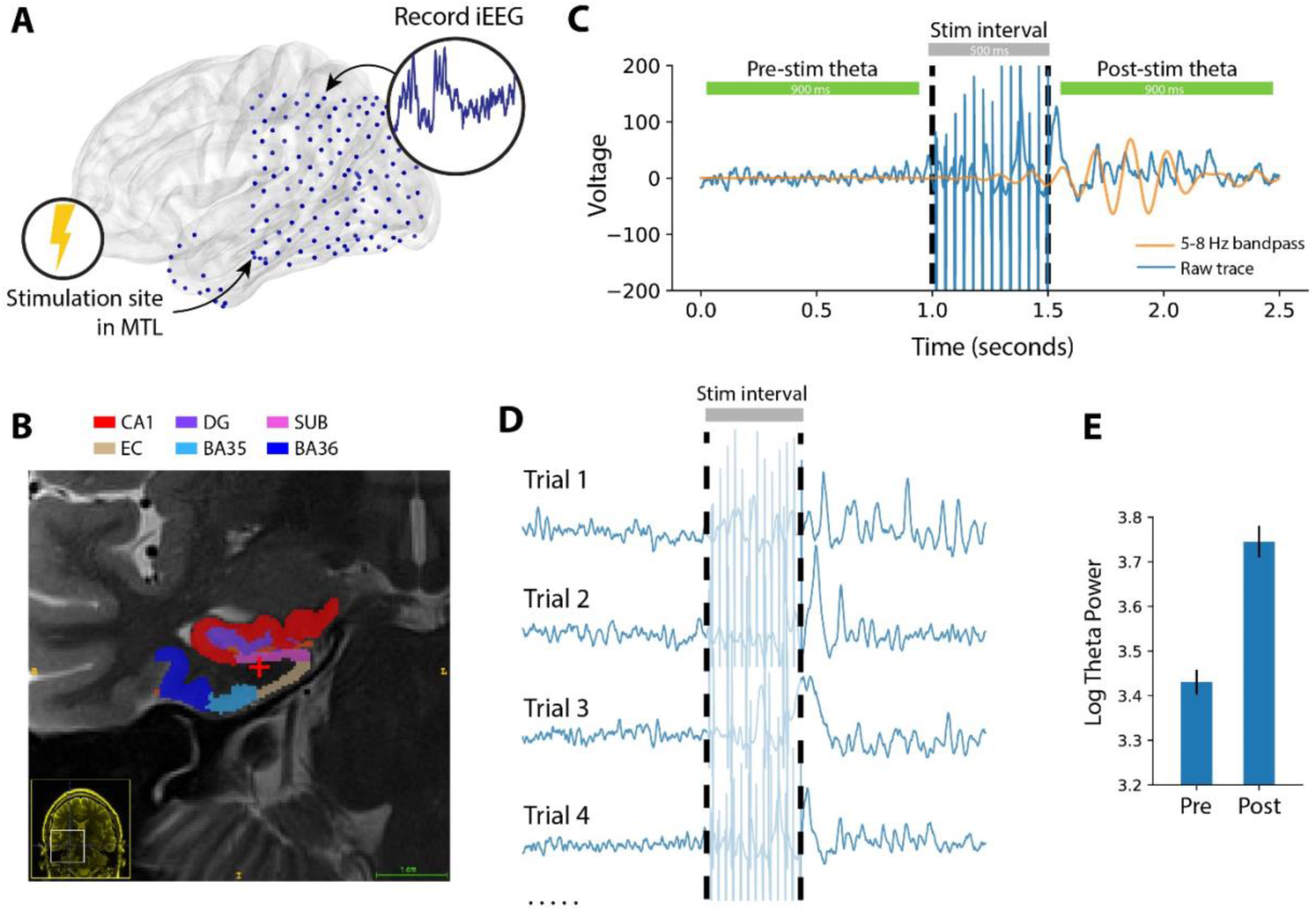
Comparison of pre- vs. post-stimulation theta (5-8 Hz) power in an example subject. **(A)** Each of 26 subjects received a series of 500 ms bipolar stimulation events, at 1-7 sites within the MTL; an example subject schematic is shown here. **(B)** T2 MRI and MTL subregion segmentation for an example subject. Stimulation location, in white matter, is indicated at the red cross. See Supplemental Figure 1 for subregion labels. **(C)** Using the multitaper method, theta power (5-8 Hz) was measured in 900 ms windows preceding and following each stimulation event, with 50 ms buffers before and after stimulation. In an example stimulation event, the 5-8 Hz bandpass signal (orange) is overlayed on the raw bipolar signal (blue), to emphasize a change in pre- vs. post-stimulation theta power. **(D)** Theta power is extracted in the pre- and post-stimulation intervals for at least 240 events (“trials”) per stimulation site. **(E)** The log-transformed theta power is aggregated for all pre- and post-stimulation intervals separately, for later statistical comparison (Fig. 2).

**Figure 2.**
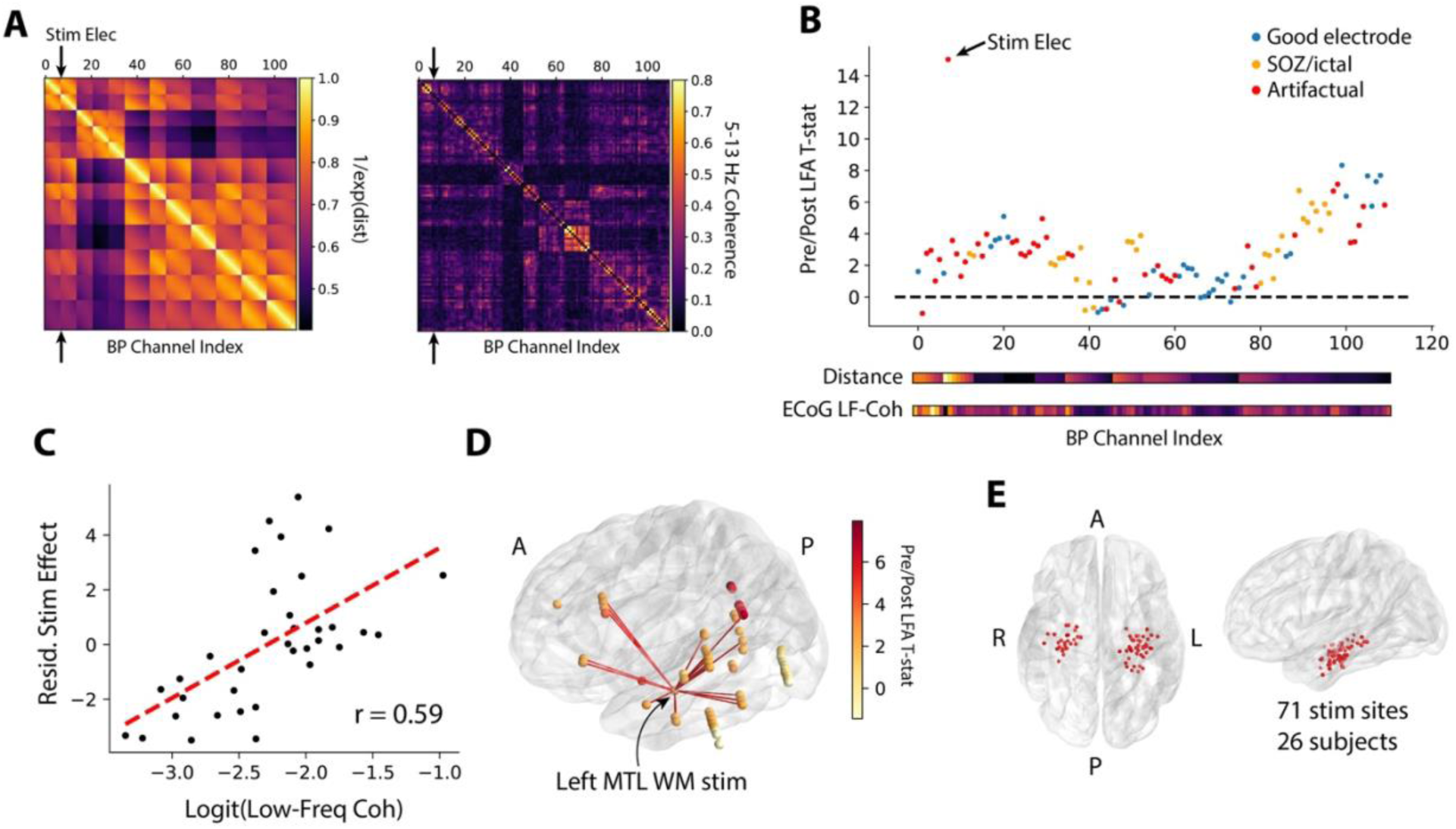
Method for determining theta modulation index (TMI). **(A)** For each subject, Euclidean distances (left matrix) and functional connectivity (right matrix) are measured for all possible electrode pairs. Distances are linearized as e^-(distance)^, with 1.0 representing no separation between two electrodes. Functional connectivity is the averaged 5-13 Hz multitaper coherence in 1-second windows extracted from a baseline period. **(B)** Pre- and post-stimulation theta power (Fig. 1C) is compared with a paired t-test to generate a *t*-statistic for each electrode. Electrodes are excluded from analysis if they exhibited significant post-stimulation artifact (red, see Methods for details) or were placed in the seizure onset zone or exhibit high inter-ictal spiking (orange). **(C)** Multiple linear regression is used to correlate the functional connectivity (between a recording electrode and the stimulation electrode) with the power *t*-statistic, independent of distance. To demonstrate this, the distance-residualized *t*-statistic (“Stim Effect”) is plotted against functional connectivity in the example subject. The z-scored version of this correlation is referred to as the “Theta Modulation Index,” or TMI. **(D)** Rendering of the power *t*-statistic as color on each electrode in the example subject, plotted with the top 10% of functional connections to the stim electrode (red lines). **(E)** Anatomical distribution of all MTL stimulation sites in the 26-subject dataset.

For each stimulation trial, we computed theta power (5-8 Hz) in 900 ms windows before and after each 500 ms stimulation event, and compared the pre- vs. post-stimulation power across all trials with a paired *t*-test (Figure 1D). Next, we used linear regression to correlate the strength of a stimulation site’s network connectivity to a recording electrode with the power t-statistic at that electrode (Figure 2A-D). We included absolute distance as a factor in our regression, to only consider how connectivity relates to stimulation beyond the brain’s tendency to densely connect nearby regions ^20^. The result is a model coefficient that indicates, independent of distance, the degree to which functional connectivity predicts stimulation-induced change in theta power at a recording site. The regression was repeated using permuted connectivity/evoked power relationships to generate a null distribution of model coefficients against which the true coefficient is compared. We refer to the resulting z-score as the “theta modulation index,” or TMI. High TMIs indicate functional network connectivity predicts observable stimulation-related change in theta power at distant sites.

### TMI is correlated with proximity to white matter

At a group level of stimulation sites, TMI was significantly greater than zero (1-sample *t*-test, *t*(71) = 4.0, *P* = 0.0002; Figure 3A), indicating that stimulation in the MTL tends to evoke network-driven change in theta power in distant regions. However, we noted substantial heterogeneity between stimulation sites, with some showing little or no ability to modulate network-wide theta activity, as reflected by TMIs near zero. To explain this heterogeneity, we hypothesized that, as earlier work demonstrated ^11,21,22^, structural connections (i.e. white matter tracts) may be key to the propagation of electrical stimulation throughout the brain.

**Figure 3.**
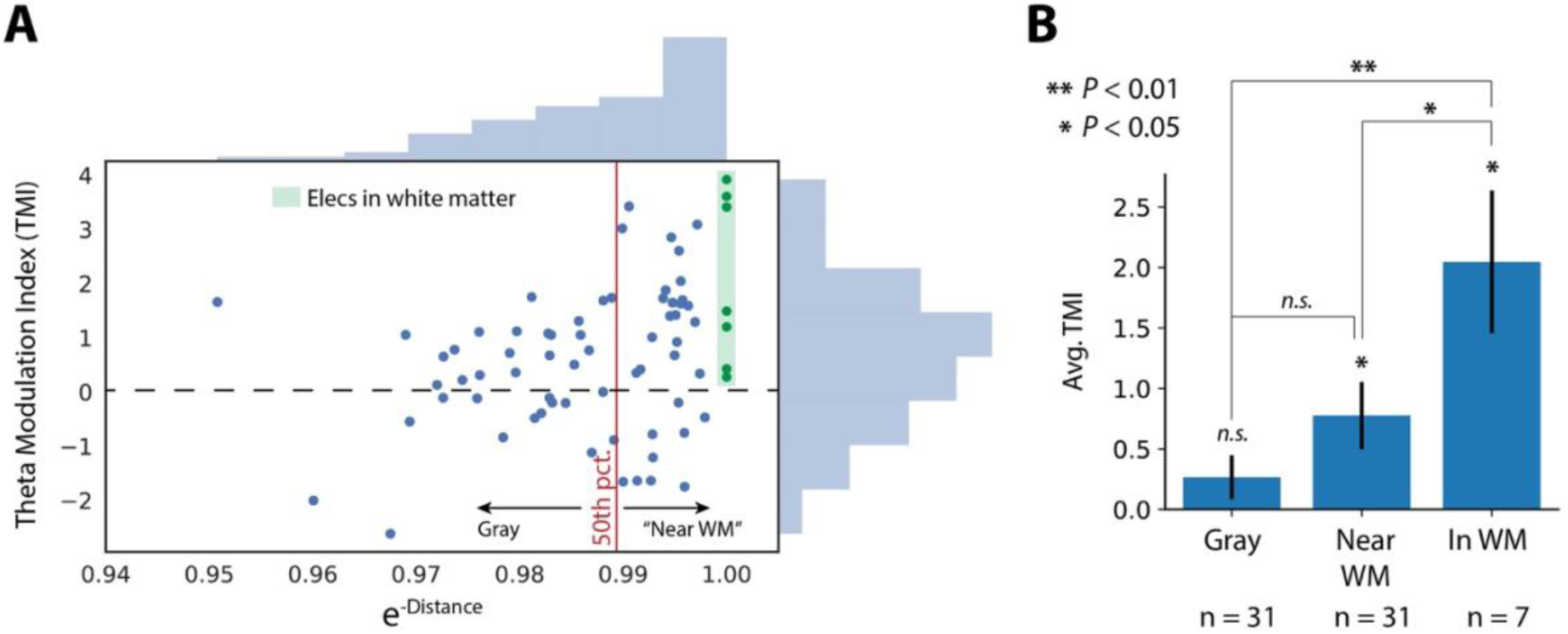
Proximity to white matter predicts TMI. **(A)** Correlation between a stimulation site’s distance from nearest white matter with the site’s TMI. The 50^th^ percentile of white matter distances divides sites classified as “gray matter” versus “near white matter.” Stimulated contacts in white matter are highlighted in green. See Supplemental Figure 2 for the Pearson correlation of these data (*r* = 0.33, *P* = 0.005). **(B)** TMI increases with closeness to white matter, as determined by a permutation test (*P* < 0.001, see Methods) and by noting that TMIs for sites in or near white matter are significantly greater than zero (1-sample t-test, *P* < 0.05) while gray matter sites are not (*P* = 0.15). Electrodes placed in white matter have greater TMIs than electrodes near white matter (2-sample t-test, *P* < 0.05) or gray matter (*P* < 0.01). Error bars show +/- 1 SEM; * *P* < 0.05; ** *P* < 0.01.

To test whether structural connections play a role in stimulation propagation, we asked whether TMI was correlated with the proximity of a stimulation site to white matter. If these measures are correlated, it would indicate that functional connectivity is predictive of physiology only insofar as white matter tracts are accessible. We binned stimulation sites according to whether they were placed in gray matter (n = 32, lower 50^th^ percentile of distances to white matter), near white matter (n = 33, upper 50^th^ percentile of distances to white matter), or within white matter (n = 7, manually identified by a neuroradiologist; Figure 3A; see Supplementary Figure 1 for anatomical placement of each white matter target). We found that TMI was significantly increasing with white matter placement, relative to a permuted distribution (permuted *P* < 0.001; Figure 3B). The TMI for gray matter sites was not significantly different than zero (1-sample *t*-test, *t*(31) = 1.4, *P* = 0.18), while TMI for sites near or in white matter was significant (*P* < 0.05). This relationship holds in a Pearson correlation agnostic to any electrode categorization (*r* = 0.33, *P* = 0.005; Supplementary Figure 2). This finding does not mean gray matter stimulation fails to induce theta activity, but it does suggest gray matter stimulation may result in theta activity that is uncorrelated with functional connectivity to remote sites.

Taken together, these results show that direct electrical stimulation of the MTL can induce spectral power changes across a distributed network of regions, particularly if stimulation occurs in or proximal to white matter. When this occurred, we discovered that functional low-frequency coherence is predictive of where stimulation-related modulations in theta power are observed.

### Network properties of MTL stimulation

Having shown that stimulation in or near white matter sites induces distributed changes in theta power, we next sought to characterize the directionality of change. Specifically, high TMIs could be caused by increases in theta power at electrodes with strong functional connectivity to the stimulation target, or decreases in theta power at electrodes with weak connectivity to the stimulation target. To distinguish between these possibilities, we further examined theta power changes in detail among the 16 stimulation sites that exhibited significant (*P* < 0.05) TMI (see Supplementary Table 1 for statistics and anatomical placement of each significant site). In this subset, we measured the average pre- vs. post-stimulation theta power at the five electrodes with the strongest functional connectivity to the stimulation site (controlled for distance), and the five electrodes with the weakest functional connectivity. At strongly-connected sites, theta power change was significantly positive (1-sample *t*-test, *t*(15) = 5.6, *P* = 4.0 × 10^−5^) and significantly greater than power change at weakly-connected sites (paired *t*-test, *t*(15) = 6.03, *P* = 1.7 × 10^−5^; Figure 4B). No significant power change was observed at sites with weak functional connectivity (1-sample *t*-test, *t*(15) = 1.5, *P* = 0.15). Notably, we observed that of the 16 significant sites analyzed here, 15 were placed in or near white matter. We conclude that stimulation causes increased theta power at strongly-connected sites and little to no change in power at weakly-connected sites.

**Figure 4.**
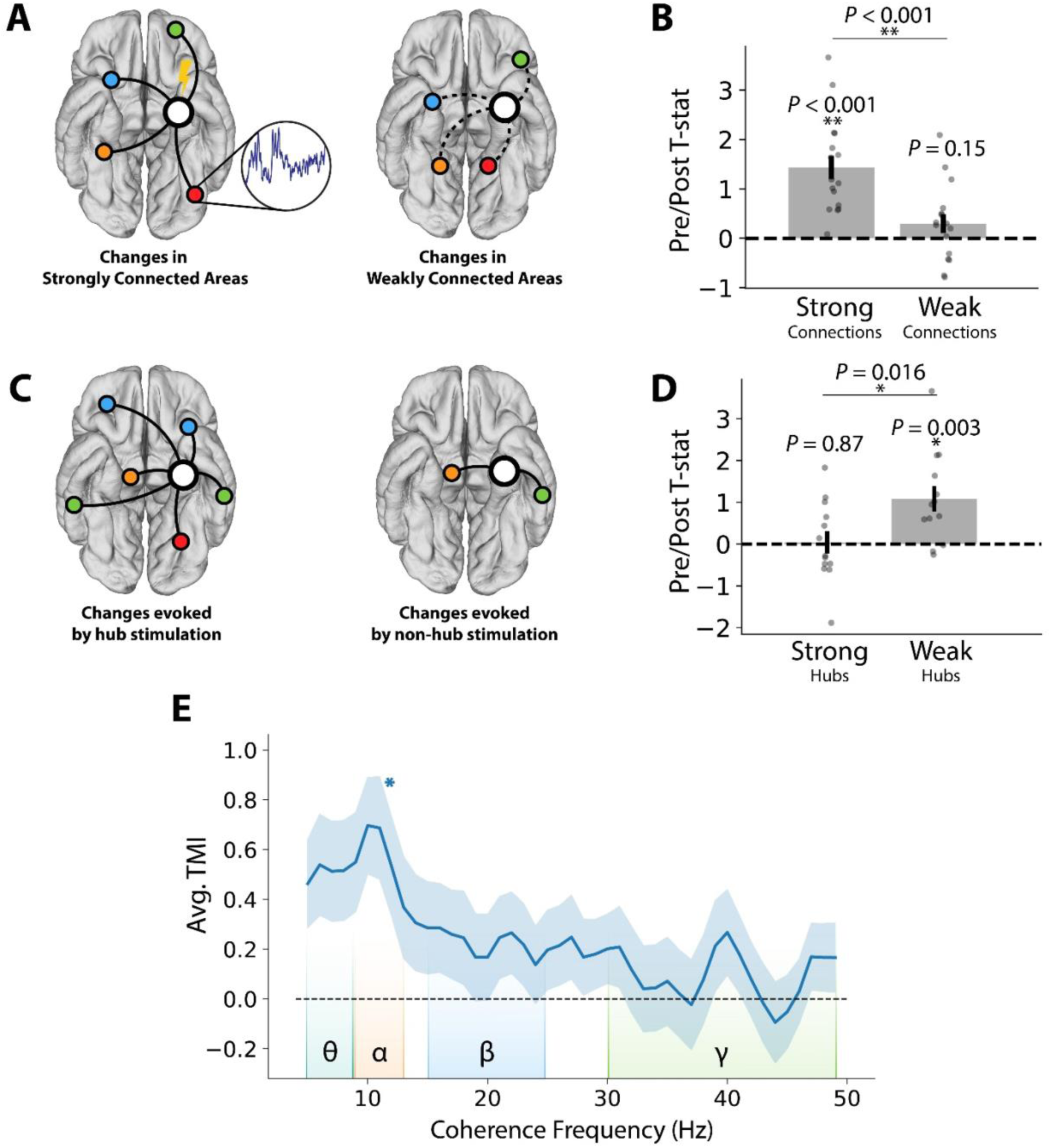
Network properties of stimulation-induced theta. **(A)** Schematic of a stimulation site and its most strongly-connected areas (left) or weakly-connected areas (right). **(B)** For each of 16 stimulation sites with significant TMI (*P* < 0.05), the average post- vs. pre-stimulation theta T-statistic is computed for the five strongest-connected electrodes and the five weakest-connected electrodes (controlled for distance). Changes at strongly-connected recording sites are significantly greater than changes at weakly-connected sites (paired *t*-test, *t*(15) = 6.03, *P* = 1.7 × 10^−5^). **(C)** Schematic of a hub-like stimulation site (left) and a non-hub stimulation site (right). Hub scores are calculated as the node strength, or average of all connection weights to a given electrode. **(D)** For each of 40 stimulation sites in or near white matter, the average post- vs. pre-stimulation theta T-statistic is computed for the five strongest-connected recording electrodes. Stimulation of a weak hub (lower tercile of hub scores, n = 13) yields significantly greater change in connected regions than stimulation of a strong hub (upper tercile of hub scores, n = 14) (2-sample *t*-test, *P* = 0.016). **(E)** Average TMI across all in or near-white matter stimulation sites, as a function of functional connectivity frequency. TMI is greatest for networks constructed from theta or alpha coherence (5-13 Hz). Corrected for multiple comparisons across all frequencies, TMI is significantly greater than zero at 11 Hz. Error bars show +/- 1 SEM; * *P* < 0.05; ** *P* < 0.001.

Principles of network control theory suggest a relation between the connectivity profile – or network topology – of a stimulation site and the ensuing change in brain activity. Network “hubs,” or regions with strong connectivity to the rest of the brain, are generally less capable of modulating the brain’s overall state versus non-hubs, or regions with strong connections to only a few areas ^14,23^. To directly test this hypothesis, we asked whether stimulation-induced theta power correlated with the functional “hubness” of a stimulation site. We again took our measure of stimulation-induced activity to be the theta power change at the 5 recording sites with the strongest functional connectivity to the stimulation site, and tested this metric against the node strength of a stimulation site, an indicator of hubness (for this analysis, we considered all stimulation sites in or near white matter; n = 40). When weak hubs (lower tercile of hub scores; n = 13) were stimulated, power change at connected recording sites was significantly greater than zero (1-sample t-test, *t*(12) = 3.6, *P* = 0.003), but stimulation at strong hubs (upper tercile; n = 14) evoked no significant power modulation (*t*(13) = 0.15, *P* = 0.87; Figure 4D). While counterintuitive, this result is in line with the prediction of network control theory; stimulation at a site with many connections may disperse the effect of perturbation, yielding lesser activation in downstream regions.

Our choice of low-frequency (5-13 Hz) functional connectivity as the basis for predicting distributed changes in theta power was motivated by prior studies that have shown strong, cognitively-relevant connectivity at low frequencies particularly the theta and alpha bands ^16,17,19^. However, others have noted significant inter-regional connectivity in the beta and gamma bands 24. As our study presented a unique opportunity to examine the causal nature of functional connectivity, we asked whether functional connectivity in other frequency bands is also predictive of downstream power modulations. Among all MTL electrodes placed in or near white matter (n = 40), we asked whether TMI was significant for networks constructed from any frequency to a maximum of 50 Hz. No frequencies outside the alpha/theta bands exhibited significant group-level TMIs, after correction for multiple comparisons (*P* < 0.05, Benjamini-Hochberg correction; Figure 4E). This demonstrates that functional networks constructed from high frequencies (> 13 Hz) are not predictive of stimulation-induced theta activity.

## Discussion

We set out to test a fundamentally simple hypothesis: Do functional connections in the brain predict how focal electrical stimulation flows from one region to another? Though critical to the future of brain stimulation and therapeutic development, this hypothesis has not seen rigorous testing. Prior studies indicate that connectivity plays a role in how stimulation events perturb distant brain regions ^10,12,21,22^, but fundamental assumptions of graph-theoretic models remain untested ^13^. More broadly, no prior studies have addressed whether iEEG-based functional connectivity indicates anything about causal relationships in the brain, or whether is it merely a correlative measure. In this manuscript, we specifically tested a hypothesis about the effects of stimulation on theta power, given an especially rich literature showing the cognitive relevance of theta oscillations ^25–28^. This theoretically-motivated choice also served to reduce the number of possible tests we could have run on other frequency bands implicated in cognition.

We discovered that (1) modulation of low-frequency power is correlated with functional connectivity, but only if stimulation occurred in or near white matter, (2) stronger functional connections yield greater power increases, (3) stimulation of strong functional hubs weakens downstream power changes, and (4) low-frequency functional connections are more strongly predictive of neural activity than high frequency connections. These results align with predictions offered by models of the brain as a dynamical system built on a white matter scaffold. Namely, stronger connections generally yield bigger evoked changes, but too many strong connections can dilute the effect of stimulation – perhaps by dispersing stimulation energy across many regions.

The meaning of functional connectivity is a subject of considerable debate. Correlated activity between two parts of the brain may reflect direct connection between the two, an indirect connection through a third region, or the activity of a third region independently driving activity in each ^29^. Though most neuroscientists are aware of such limitations, functional connectivity is often implicitly treated as a measure of causality nonetheless. Our use of targeted stimulation allowed us to test whether this implicit assumption is true. Our results generally support the idea that functional connectivity indicates causality; when stimulation occurs in or near white matter, we could predict where power changes would occur based on distance-independent measures of low-frequency functional connections. This finding aligns with observations that intrinsic functional connectivity in MRI is constrained by white matter anatomy ^30^. However, substantial variance in power modulation remained unexplained by connectivity, and we also showed that propagation of gray matter stimulation - still rich with functional connections - cannot be predicted in the same way.

This study faces several limitations. First, we only assessed stimulation in the MTL, which has a distinct architecture that may affect how stimulation propagates to other regions – the effects of stimulation at the cortical surface could differ markedly. High-resolution diffusion tractography would be needed to make strong claims about which MTL white matter tracts are accessed by stimulation. Second, though our hypothesis about theta was theoretically grounded, this choice leaves open the question as to whether higher frequency activity is also affected by stimulation – though theta power may be a useful measure of how stimulation alters neural excitability ^31–33^, changes in population spiking would be better captured by high-frequency activity (e.g. > 60 Hz) ^34,35^.

In this study we solely analyze stimulation through the lens of changes in brain physiology. However, with an eye towards the eventual therapeutic use of stimulation, the results here bridge prior studies of stimulation and behavior with underlying neural mechanisms. A recent study reported decreases in episodic memory performance during stimulation at certain times, associated with increases in cortical theta power ^3^. Additionally, memory performance was noted to increase with theta-burst stimulation of the perforant path, a major white matter tract of the MTL ^4^. Deep brain stimulation targeted to white matter tracts has also been shown to improve outcomes in treatment-resistant depression ^9^. Collectively, these findings are supported by the results here - white matter stimulation appears to evoke remote increases in neural activity. Few studies have deeply examined stimulation-induced changes in physiology with behavioral enhancement, though our approach outlined here enables us to do exactly that in future work.

Here we demonstrated that functional connections in the human brain inform how stimulation evokes remote changes in neural activity. This is powerful new evidence that, even in the absence of knowledge about an individual’s structural connectome, functional connectivity reflects causality in the brain – a finding with significant implications for how neuroscientists interpret inter-regional correlations of neural activity. Furthermore, by showing that stimulation-evoked changes interact with the functional hubness of a targeted site, we provided critical, empirical evidence that network control theory can model real-world brain dynamics.

## Supplementary Information

is linked to the online version of the paper at www.nature.com.

## Acknowledgements

We thank Blackrock Microsystems for providing neural recording equipment. This work was supported by the DARPA Restoring Active Memory (RAM) program (Cooperative Agreement N66001-14-2-4032), as well as National Institutes of Health grant MH55687 and T32NS091006. We are indebted to all patients who have selflessly volunteered their time to participate in our study. The views, opinions, and/or findings contained in this material are those of the authors and should not be interpreted as representing the official views or policies of the Department of Defense or the U.S. Government. We also thank Drs. Youssef Ezzyat, Christoph Weidemann, Nora Herweg, Danielle Bassett, and Geoffrey Aguirre for providing valuable feedback on this work.

## Author Contributions

E.S., M.J.K., and D.S.R. designed the study; E.S. conceived, planned, and executed all data analyses, J.K. analyzed data, and E.S. wrote the paper. J.S., R. Gorniak, S. Das. performed anatomical localization of depth electrodes. M.S., G.W., B.L., R. Gross., B.J., C.I., K.Z., S. Sheth, and S. Seger recruited subjects, collected data, and performed clinical duties associated with data collection including neurosurgical procedures or patient monitoring.

## Data Availability

Raw electrophysiogical data used in this study is freely available at http://memory.psych.upenn.edu/Electrophysiological_Data

## Methods

### Participants

Twenty-six patients with medication-resistant epilepsy underwent a surgical procedure to implant subdural platinum recording contacts on the cortical surface and within brain parenchyma. Contacts were placed so as to best localize epileptic regions. Data reported were collected at 8 hospitals over 4 years (2015-2018): Thomas Jefferson University Hospital (Philadelphia, PA), University of Texas Southwestern Medical Center (Dallas, TX), Emory University Hospital (Atlanta, GA), Dartmouth-Hitchcock Medical Center (Lebanon, NH), Hospital of the University of Pennsylvania (Philadelphia, PA), Mayo Clinic (Rochester, MN), National Institutes of Health (Bethesda, MD), and Columbia University Hospital (New York, NY). Prior to data collection, our research protocol was approved by the Institutional Review Board at participating hospitals, and informed consent was obtained from each participant.

### Electrocorticographic recordings

iEEG signal was recorded using depth electrodes (contacts spaced 3.5-10 mm apart) using recording systems at each clinical site. iEEG systems included DeltaMed XlTek (Natus), Grass Telefactor, and Nihon-Kohden EEG systems. Signals were sampled at 500, 1000, or 1600 Hz, depending on hardware restrictions and considerations of clinical application. Signals recorded at individual electrodes were first referenced to a common contact placed intracranially, on the scalp, or mastoid process. To eliminate potentially confounding large-scale artifacts and noise on the reference channel, we next re-referenced the data using a bipolar montage. Channels exhibiting highly non-physiologic signal due to damage or misplacement were excluded prior to re-referencing. The resulting bipolar timeseries was treated as a virtual electrode and used in all subsequent analysis.

### Anatomical localization

To precisely localize MTL depth electrodes, hippocampal subfields and MTL cortices were automatically labeled in a pre-implant, T2-weighted MRI using the automatic segmentation of hippocampal subfields (ASHS) multi-atlas segmentation method ^36^. Post-implant CT images were coregistered with presurgical T1 and T2 weighted structural scans with Advanced Normalization Tools ^37^. MTL depth electrodes that were visible on CT scans were then localized within MTL subregions (including white matter) by neuroradiologists with expertise in MTL anatomy. All localizations in this manuscript refer to the bipolar midpoint of two recording contacts or the anode/cathode stimulation contacts.

### Functional connectivity estimation

To obtain coherence values between electrode pairs, we used the MNE Python software package ^38^, a collection of tools and processing pipelines for analyzing EEG data. The coherence (*C_xy_*) between two signals is the normalized cross-spectral density (Equation 1); this can be thought of as the consistency of phase differences between signals at two electrodes, weighted by the correlated change in spectral power at both sites.

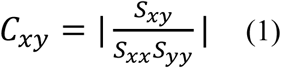

Where *S_xy_* is the cross-spectral density between signals at electrodes *x* and *y*; *S_xx_* and *S_yy_* are the auto-spectral densities at each electrode. Consistent with other studies of EEG coherence ^39,40^, we used the multitaper method to estimate spectral density. We used a time-bandwidth product of 4 and a maximum of 8 tapers (tapers with spectral energy less than 0.9 were removed), computing coherence for frequencies between 4-50 Hz, avoiding the 60 Hz frequency range that may be contaminated by line noise. Inter-electrode coherences were computed for a series of 1-second windows (minimum of 10 windows per subject) extracted from the baseline period of a non-stimulation task, in which subjects wait passively before beginning a verbal free-recall task. To construct the low-frequency networks used in the majority of this paper, cross-spectra were first averaged across all baseline period windows, normalized by the average power spectra, and then averaged between 5-13 Hz. For the analysis in Figure 4E, networks are constructed for each frequency between 4-50 Hz with no averaging over bands.

### Stimulation paradigm

At the start of each session, we determined the safe amplitude for stimulation using a mapping procedure in which stimulation was applied at 0.5 mA, while a neurologist monitored for afterdischarges. This procedure was repeated, incrementing the amplitude in steps of 0.5 mA, up to a maximum of 1.5 mA (chosen to be below the afterdischarge threshold and below accepted safety limits for charge density ^41^). For each stimulation session, we passed electrical current through a single pair of adjacent electrode contacts in the MTL. Stimulation was delivered using charge-balanced biphasic rectangular pulses (pulse width = 300 μs) at (10, 25, 50, 100, or 200) Hz frequency and (0.25 to 2.00) mA amplitude (0.25 mA steps) for 500 ms, with a minimum of 3 seconds between stimulation events. During a session, subjects were instructed to sit quietly and did not perform any task. An average of 2.7 stimulation sites were selected for each subject, with a minimum of 240 trials delivered for each.

In most subjects, a post-stimulation voltage deflection artifact briefly contaminates a subset of recording contacts. To identify and remove channels exhibiting this artifact, the average voltage in the 350 ms prior to stimulation is compared with a paired t-test to the average voltage in the 350 ms after stimulation, across all trials, for each channel. The same procedure is done with a levene test for different variances. Any electrode with a significantly different pre-vs.-post mean voltage or voltage variance (*P* < 0.01) is excluded from further analysis (see “Estimating theta modulation index”). On average, this procedure excludes 28% of channels. Regardless of stimulation artifact, any bipolar pair is excluded from analysis if it shares a common contact with the stimulated pair.

### Spectral power analysis

We used the multitaper method to assess spectral power in the pre- and post-stimulation intervals (−950 to −50 ms relative to stimulation onset, and +50 to +950 ms after stimulation offset; Figure 1B). We avoided the Morlet wavelet method to obviate the need for buffer periods that extend into the stimulation window. As in “Functional connectivity estimation,” we used the MNE Python software package. For each trial, theta power was taken as the average PSD from 5-8 Hz, using a time-bandwith product of 4 and excluding tapers with < 90% spectral concentration. To compute a T-statistic at each electrode, the pre- vs. post log-transformed power values were compared with a paired t-test (Fig. 1D, Fig. 2B). We avoid calculating significances for individual electrodes because sequential trials are non-independent events; T-statistics are only used for later correlation analysis (see “Theta modulation index”).

### Estimating theta modulation index

To examine the relationship between stimulation and functional connectivity, we developed an index that reflects the correlation between theta power modulation and connectivity, independent of distance. To do this, we first construct low-frequency (5-13 Hz) networks as described in “Functional connectivity estimation,” and take the logit transform to linearize coherence values that fall between 0 and 1. We also construct adjacency matrices that reflect the normalized Euclidean distance between all possible pairs of electrodes (Fig. 2A), and linearize the distances by taking the reciprocal of their exponential (i.e. a Euclidean distance of zero would correspond to 1.0). For each stimulated electrode, we take that electrode’s distance and connectivity to all other electrodes as predictors of the theta power t-statistic (see “Spectral power analysis) in a multiple linear regression. This controls for the effect of distance from a stimulation target, which is correlated with power and functional connectivity. Next, we permute the order of the predictors 1000 times and re-run the regression for each. The true coefficient for functional connectivity is compared to the distribution of null coefficients to obtain a z-score and p-value for each stimulation site. The z-score is referred to as the theta modulation index, or TMI.

Prior to computing TMI, we excluded electrodes placed in the seizure onset zone or exhibiting significant inter-ictal spiking, as determined by a clinician. Electrodes with high post-stimulation artifact (see “Stimulation paradigm”), and stimulated electrodes themselves, were also excluded. Subjects were discarded if less than 10 electrodes remained after all exclusions.

To analyze the relationship between TMI and white matter category (Fig. 3), we first binned electrodes according to their distance from nearest white matter. Distance were measured as the linearized Euclidean distance from a stimulation electrode (i.e. bipolar midpoint of the anode/cathode) to the nearest vertex of that subject’s Freesurfer white matter segmentation ^42^ based on T1 MRI. The 50^th^ percentile of white matter distances marked the division between stimulation electrodes categorized as “near” white matter versus in gray matter. Seven stimulation electrodes were identified by expert neuroradiologists as being placed within white matter (see Supplementary Figure 2 for exact placements). To ask whether TMI increases with white matter category, permuted the white matter labels for each electrode 1000 times and took the minimum T-statistic between gray vs. near and near vs. in categories at each permutation. We then compared the minimum T-statistic in the true data to the distribution of null statistics to generate a p-value.

### Network properties of stimulation

To determine how the network structure of a stimulation site affected downstream alterations in theta power (Fig. 4), we first analyzed the relationship between pre- vs. post-stimulation theta power and the strength of functional connectivity to a stimulation site (Fig. 4A-B). For each stimulation site with a significant TMI (*P* < 0.05), we ranked all other electrodes by the strength of their functional connectivity to that site, residualized on Euclidean distance (e^-dist^). We then took the average power T-statistic (see “Spectral power analysis”) across the 5 strongest-connected sites and the 5 weakest-connected sites, to assess whether theta power changes correlated with the strength of a functional connection.

To assess whether the effects of stimulation differ between hubs and non-hubs (Fig. 4C-D), we measured the node strength ^43^ for each stimulation site in or near white matter (n=38), using our low-frequency coherence networks (see “Functional connectivity estimation”). The node strength reflects the sum of all connection strengths to a given node (for this paper, we normalized node strength by the total number of possible connections for a given site, yielding strengths in the range from 0 to 1). For all stimulation sites, we binned hub scores by tercile, and took the highest tercile as “strong hubs,” the weakest tercile as “weak hubs” (n=13 for each). For stimulation at all strong and weak hubs, we took the average power T-statistic for the 5 strongest-connected electrodes. These values were used to assess whether hub stimulation tends to cause greater power changes in connected regions. The relationship between coherence frequency and theta modulation index (Figure 4E) was assessed by re-estimating the TMI (see “Estimating theta modulation index”) using spectral coherence networks observed for each frequency between 4-50 Hz, spaced by 1Hz, for all stimulation electrodes placed within or near white matter. The average TMI across sites/subjects was 1-sample *t*-tested against zero and p-values were FDR corrected for multiple comparisons (corrected *P* < 0.05). For visualization purposes only, the displayed TMI/frequency curve was smoothed with a 3-point moving average window.

